## Supplemental Information for "Divergent regulation of KCNQ1/E1 by targeted recruitment of protein kinase A to distinct sites on the channel complex"

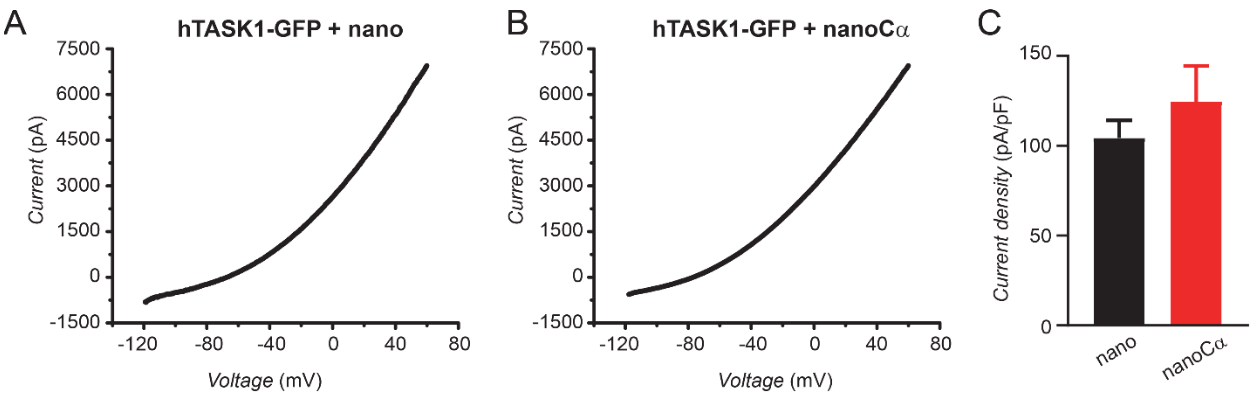


**Supplemental Figure 1: NanoCα targeted to the C-terminus of TASK1 via a GFP tag does not inhibit $K^+$ current.** (A) Exemplar current-voltage relationship elicited by a ramp stimulus (-120 to +60 mV) in a CHO cell expressing hTASK1-GFP + nano. (B) Exemplar current-voltage relationship elicited by a ramp stimulus in a CHO cell expressing hTASK1-GFP + nanoCα. (C) Population current density at 0 mV for cells expressing hTASK1-GFP with either nano (black bar, $n$ = 7) or nanoCα (red bar, $n$ = 9).

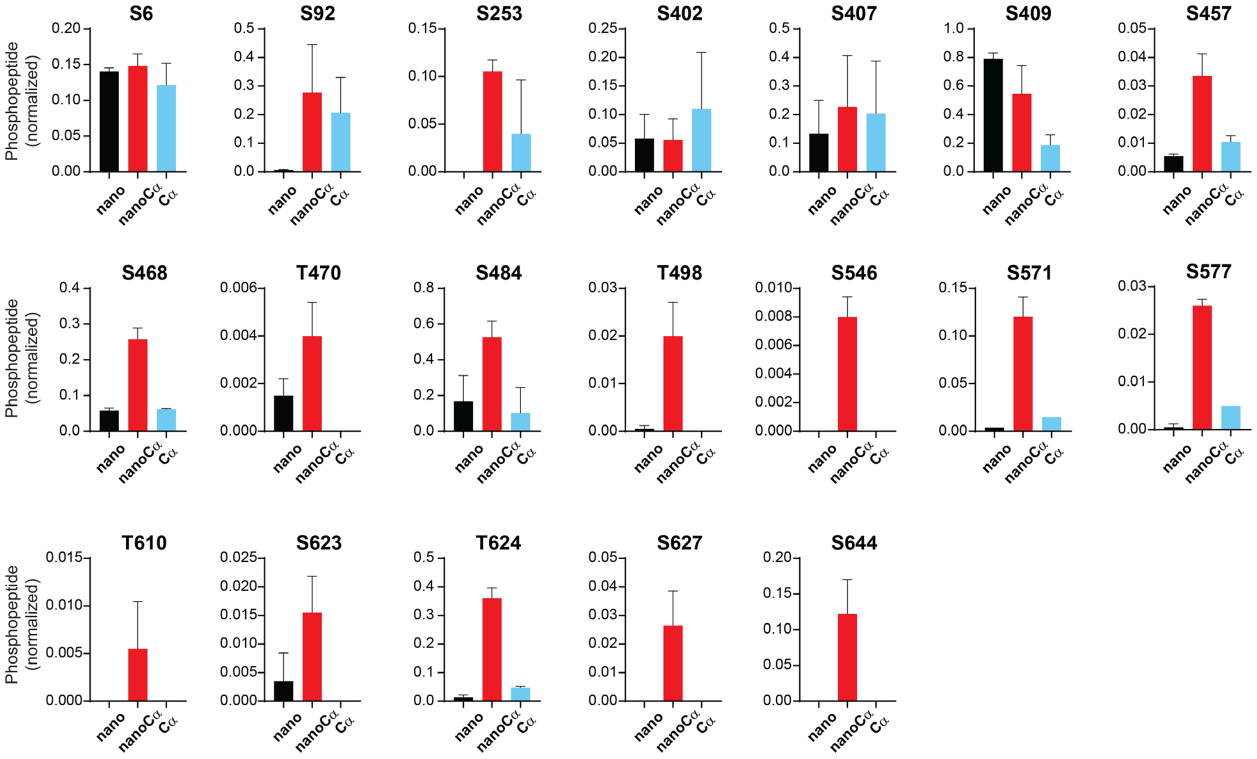


**Supplemental Figure 2:** Relative abundance of phosphorylated KCNQ1-YFP peptides identified using mass spectrometry in cells co-expressing nano (black), nanoCα (red), or free Cα (cyan).
